# New Insights into the Nature of Symbiotic Associations in Aphids: Initial Steps Involved in Aphid Gut Infection by *Serratia symbiotica* Bacteria

**DOI:** 10.1101/457945

**Authors:** Pons Inès, Renoz François, Noël Christine, Hance Thierry

## Abstract

Symbiotic microorganisms are widespread in nature and can play a major role in the ecology and evolution of animals. The *aphid-Serratia symbiotica* bacterium interaction provides a valuable model to study mechanisms behind these symbiotic associations. The recent discovery of cultivable *S. symbiotica* strains having the possibility of free-living lifestyle allowed us to simulate their environmental acquisition by aphids to examine the mechanisms involved in this infection pathway. Here, after oral ingestion, we analyzed the infection dynamic of cultivable *S. symbiotica* strains during the host’s lifetime using qPCR and fluorescence techniques and determined the immediate fitness consequences of these bacteria on their new host. We further examined the transmission behavior and phylogenetic position of cultivable strains. Usually, *S. symbiotica* are considered as maternally-transmitted bacteria living within aphid body cavity and bringing some benefits to their hosts despite their costs. Otherwise, our study revealed that cultivable *S. symbiotica* are predisposed to establish a symbiotic association with new aphid host, settling in its gut. We showed that cultivable *S. symbiotica* colonized the entire aphid digestive tract following infection, after which the bacterium multiplied exponentially during aphid development. Our results further revealed that gut colonization by the bacteria induce a fitness cost to their hosts. Nevertheless, it appeared that they also offer an immediate protection against parasitoids. Interestingly, cultivable *S. symbiotica* seem to be extracellularly transmitted, possibly through the honeydew. These findings provide new insights into the nature of symbiosis in aphids and the mechanisms underpinning these interactions.

**Importance:** For the first time, our study provides experimental data that highlight a new kind of symbiotic associations in aphids. By successfully isolating microbial symbiont from aphids and by cultivating it *in vitro* in our laboratory, we established artificial association by simulating new bacterial acquisitions involved in aphid gut infection. Our results showed the early stages involved in this route of infection. Until now, *Serratia symbiotica* is considered as a maternally-transmitted aphid endosymbiont. Nevertheless, here, we showed that our cultivable strains having an intermediate status between a strict free-living bacterium and a facultative endosymbiont, occupy and replicate in aphid gut and seem to be transmitted over generations through an environmental transmission mechanism. Moreover, they are both parasites and mutualists given the context, as many of the endosymbionts in aphids. Our findings give new perception of associations involved in aphids’ symbiosis.

## Introduction

Symbiotic associations between animals and microorganisms are ubiquitous in nature (1). These symbiotic microorganisms can profoundly exert influences on host ecology and evolution (2–4). By being a source of evolutionary innovations, some symbionts have allowed their hosts to colonize otherwise uninhabitable environments (5). This is the case for many insects that feed exclusively on nutritionally restricted diets, such as plant sap and blood, but which harbor obligate symbionts that supply essential nutrients lacking in their diets (6–8).

Obligate insect-microbe associations exhibit host-symbiont phylogenetic congruence, indicating that they are evolutionary ancient and stable associations maintained by a strict vertical transmission of symbionts from mother to progeny (9–11). Because of the long-term association with their host, genomes of these symbionts has been drastically reduced, which severely hamper their capability to survive outside their host (12). In addition to obligate microbial partners, insects can also harbor facultative symbionts which are not essential for host survival and reproduction, but which can be associated with beneficial or detrimental effects depending on the ecological or environmental context (2, 3, 13). Unlike obligate symbionts which are most of the time hosted in specialized cells called bacteriocytes, facultative partners can inhabit within gut, tissues and/or cells (6), can experience occasional horizontal transfers (14, 15) and can be acquired from environmental sources (16–18, 2). These mutualist bacteria have evolved such that they have lost the capacity to live without their hosts. This process is however poorly understood, partly because symbiosis is mainly studied using co-evolved associations rather than associations in their early stages and partly because it is difficult to cultivate bacteria that have extreme host dependence (19).

Aphids (Hemiptera: Aphididae) offer a valuable model to better explore mechanisms influencing these associations. Indeed, they harbor an obligate symbiont *Buchnera aphidicola*, which provides essential amino acids (20), is localized in specialized cells called bacteriocytes and is stably maintained by a strict vertical transmission (10, 11). Aphids can also harbor various facultative endosymbionts, whose association is more recent (21). These facultative partners are located in sheath cells, hemolymph and secondary bacteriocytes of aphids (22) and are vertically transmitted, although occasional horizontal transfers could also occur (15). Previous studies highlighted that these symbionts can confer an array of benefits to their host in certain ecological or environmental conditions (4, 23, 24). However, they can also impose a fitness cost on their host (21, 25).

Among these facultative symbionts, *S. symbiotica* is one of the most common in natural aphid populations (19). In the literature, it is known for being an aphid endosymbiont, with different strains exhibiting distinct biological characteristics (26). In the subfamily *Lachninae, S. symbiotica* includes strains that supplement the metabolic abilities of *B. aphidicola* for tryptophan synthesis and has been described as a co-obligate partner (27, 28) whereas in *Acyrthosiphon pisum*, it includes facultative endosymbionts that can be associated with heat stress tolerance and parasitoid resistance (29, 30). In addition, some strains have been isolated from aphids in the *Aphis* genus and cultivated freely in axenic pure medium without insect cell lines and FBS (fetal bovine serum) (31, 32). The associated biological effects of these cultivable strains that have potential free-living capacity remain unknown for aphids as well as the way they infect them. Furthermore, the tissue tropism has not really been investigated, but a new field study revealed that these cultivable strains were very phylogenetically close to *S. symbiotica* strains residing in the gut of field-collected aphids (mostly the *Aphis* genus) (33). These discoveries suggest that the cultivable strains can be derived from the gut of aphids, which raise the question of the actual endosymbiotic status generally attributed to *S. symbiotica* and open up new perspectives to study the nature of associations involved in aphids’ symbiosis.

A recent study showed that the cultivable strain CWBI-2.3^T^ (34) has a larger genome size, as well as more coding DNA sequences (CDSs) and, transfer and ribosomal RNAs compared to other sequenced endosymbiont *S. symbiotica* strains, signifying genome characteristics are intermediate between a free-living bacterium and a facultative endosymbiont (35). Other studies showed that the strain also conserved a larger repertoire of genes related to metabolism and an array of molecular tools facilitating the invasion of tissues in new hosts such as virulence factors (34, 36). The possibility to cultivate these strains, their potential independence with respect to their hosts as well as the genomic features suggest that they are probably in the beginning of their symbiotic life with aphids (37) and may represent a kind of missing link in the evolution from free-living towards host-dependent mutualistic symbiont (34). These cultivable bacteria are of great interest because they offer a unique opportunity to explore symbiotic associations in their early stages.

Here, we investigated the initial steps involved in aphid gut infection by the cultivable *S. symbiotica* strains to apprehend mechanisms underpinning this new association. We examined the infection dynamics of the bacteria during the host’s lifetime, using fluorescence and qPCR techniques. We then analyzed the immediate host fitness costs and benefits resulting from that formed interaction. We have also studied the transmission mode of the bacteria and assessed the relatedness of these cultivable strains. Our results revealed that cultivable *S. symbiotica* rapidly spread through the aphid gut inducing a clear fitness cost to their host but offering in return immediate fitness benefits under environmental stress. They also showed that cultivable *S. symbiotica* seem to be transmitted over generations through an external transmission mechanism. These new observations suggest that gut-associated bacteria are likely a new type of interaction in aphids and/or represent a possible evolutionary step towards an endosymbiotic association with aphids.

## Results

### Infection dynamic of cultivable *S. symbiotica* during aphid development

Fluorescence analyses revealed that cultivable *S. symbiotica* S1+ (in red) multiplies and persists through the whole digestive tract of infected aphids (Fig. 1). In the first infection step (0 days post-ingestion), *S. symbiotica* formed an aggregate in the stomach (Fig. 1B). Two days after infection, bacterial cells migrated into the intestine (Fig. 1C), after which the population of bacteria increased and dispersed into the whole gut 5 and 10 days post-infection (Fig. 1D and Fig. E). After 15 days of infection, bacteria continued to multiply strongly and further diffused along the digestive tract to reach the hindgut (Figs. 1F-H). A large mass can be observed that seems to correspond to digestive tract swelling. The negative control shows only the obligatory symbiont *B. aphidicola* in bacteriocytes (Fig. S1, Supporting information).

**FIG 1.**
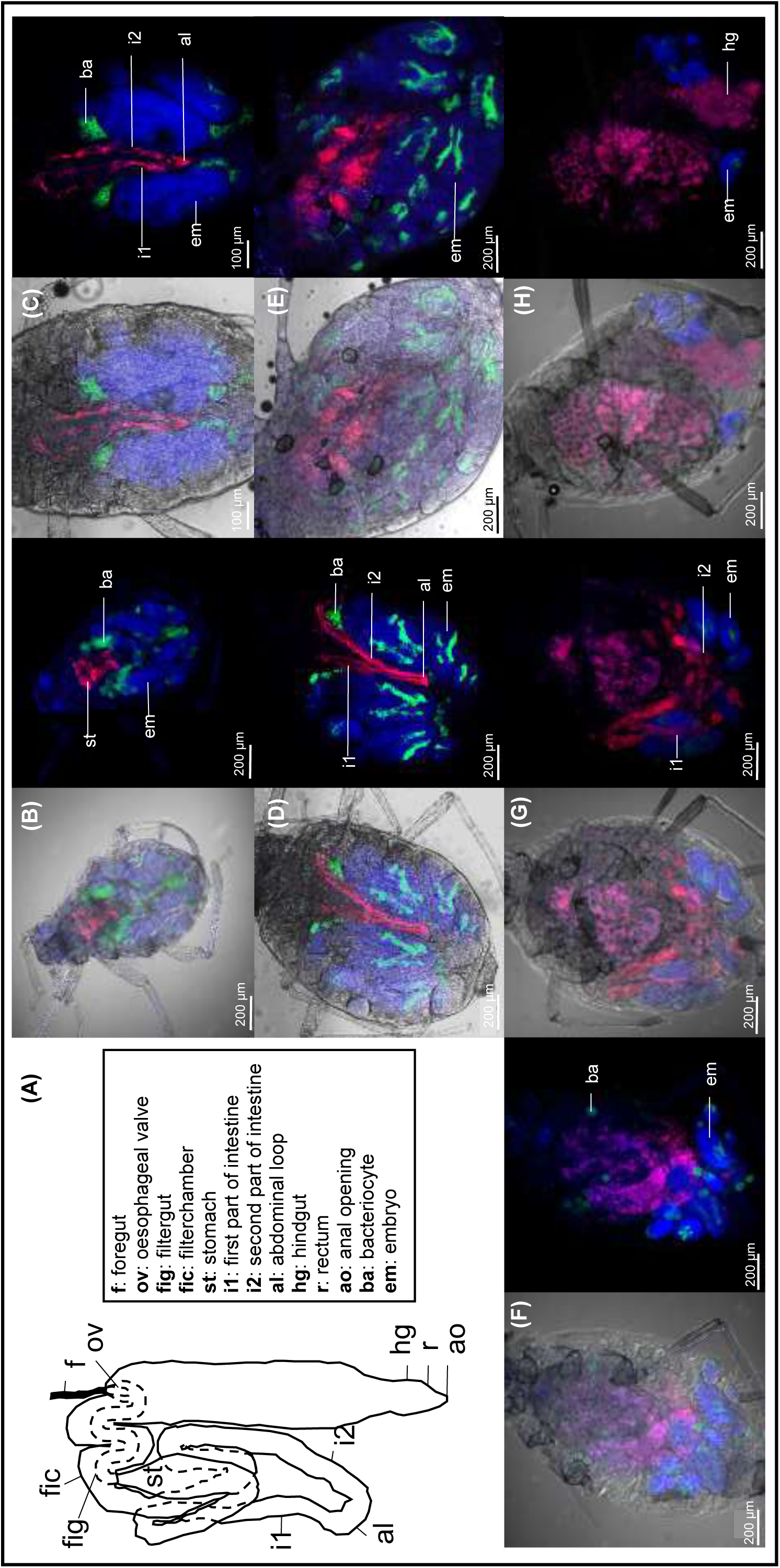
Infection processes of cultivable *S. symbiotica* CWBI-2.3^T^ (**S1+**) in *A. fabae* visualized by fluorescence *in situ* hybridization. **(A)** Semi-schematic representation of the digestive system of aphids inspired by (42), **(B)** Ingestion of *S. symbiotica* after feeding of contaminated diet (5 days-old), **(C)** 2 days post-infection (7 days-old), **(D)** 5 days post-infection (10 days-old), **(E)** 10 days post-infection (15 days-old), **(F)** 15 days post-infection (20 days-old), **(G)** 20 days post-infection (25 days-old), and **(H)** 30 days post-infection (35 days-old). For each letter corresponds two photos: on the right is without bright field and on the left is with bright field. Red Cy3 signals correspond to *S. symbiotica*, green Cys5 signals correspond to *B. aphidicola* and blue SYTOX green signals correspond to host tissues.

Quantitative PCR confirmed the infection dynamic of cultivable bacteria quantitatively. The density of *S. symbiotica* increased exponentially with aphid age (Fig. 2). Copy numbers of the *A. fabae EF1α* gene increased until 10 days post-infection (15 days of age) and then declined very slowly with aphid age (Fig. S2A, Supporting information). During oral ingestion, aphids acquired about 10^4^ copy numbers of *S. symbiotica dnaK* gene and the copy numbers of this gene increased rapidly up to 5 days post-infection (10 days of age). Then, the gene copy numbers continued to increase more slowly with aphid age to reach about 10^7^ copy numbers of *S. symbiotica dnak* gene (30 days of age) (Fig. S2B, Supporting information).

**FIG 2.**
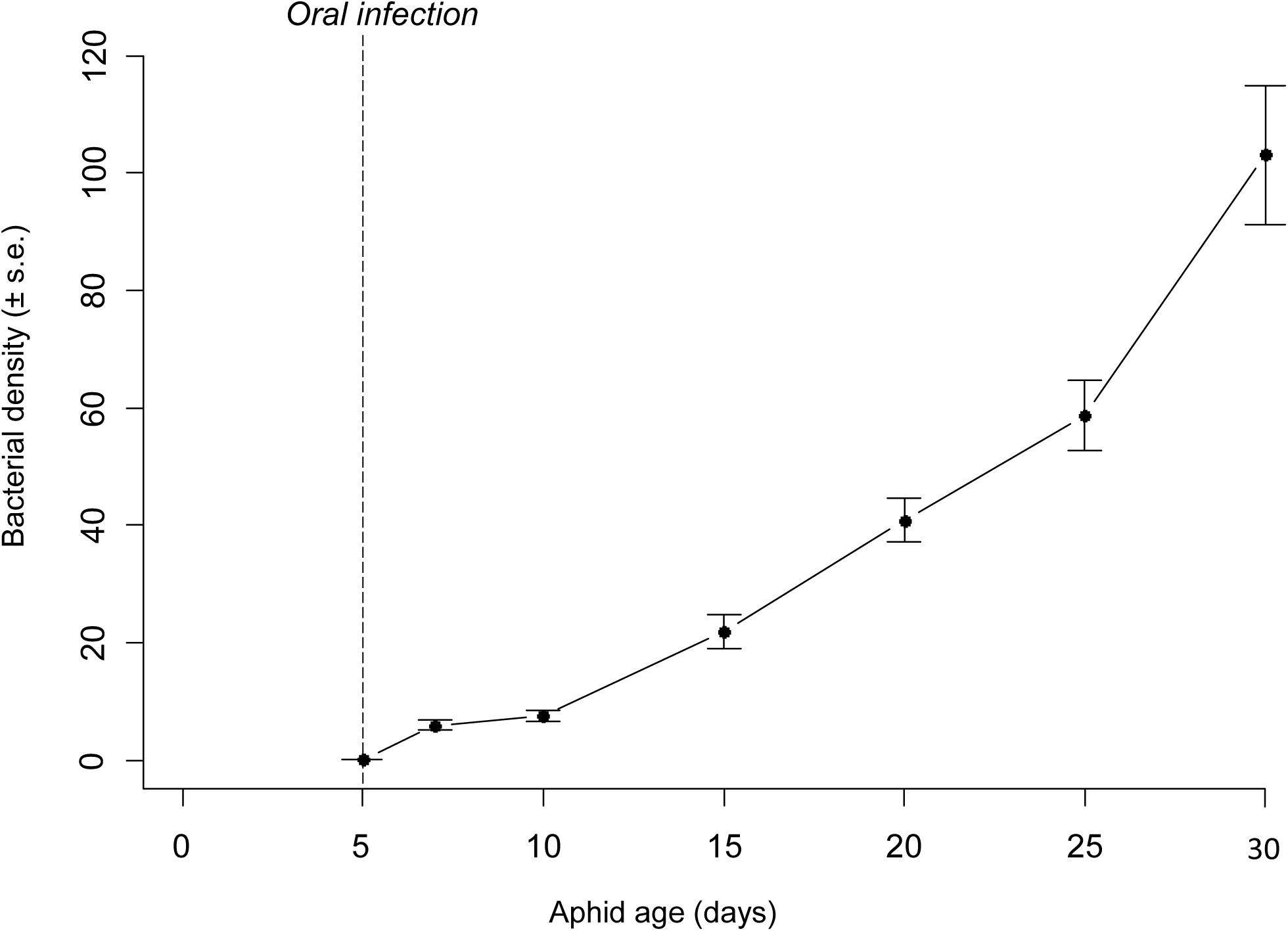
Population growth of cultivable *S. symbiotica* CWBI-2.3^T^ (**S1+**) during aphid development, estimated by quantitative PCR. Bacterial density is expressed in *S. symbiotica dnaK* gene copy numbers divided by *A. fabae EF1α* gene copy numbers (to correct for aphid body size). Error bars depict the standard error.

### Costs associated with cultivable *S. symbiotica*

All cultivable *S. symbiotica* strains had a significant negative effect on the total number of offspring (GLM, F = 20.35, df = 3, p < 0.001), adult mass (LM, F = 5.39, df = 3, p = 0.003) and duration of reproduction (GLM, F = 19.98, df = 3, p < 0.001) of *A. fabae.* Fecundity is lowered by about half for aphids infected with the three strains of *S. symbiotica* compared to uninfected aphids (Fig. 4E). Fecundity was, however, less affected in aphids harboring the **S1+** strain compared to aphids harboring the other strains (**S2+** and **S3+**). Adult mass and duration of reproduction strongly declined when aphids were infected (Figs. 4C&F). Infections also significantly affected survival of host aphids (Cox’s model, χ^2^ = 68.7, df = 3, p < 0.001). A reduction of about 20 days in longevity of infected aphids was observed compared to uninfected aphids (Fig. 4D, Table 1). Nevertheless, infection with cultivable *S. symbiotica* strains did not significantly affect nymph survival rate (GLM, χ^2^ = 159, df = 3, p = 0.058; Fig 4B) nor development time (GLM, F = 0.25, df = 3, p = 0.86; Fig 4A) of aphids.

**FIG 4.**
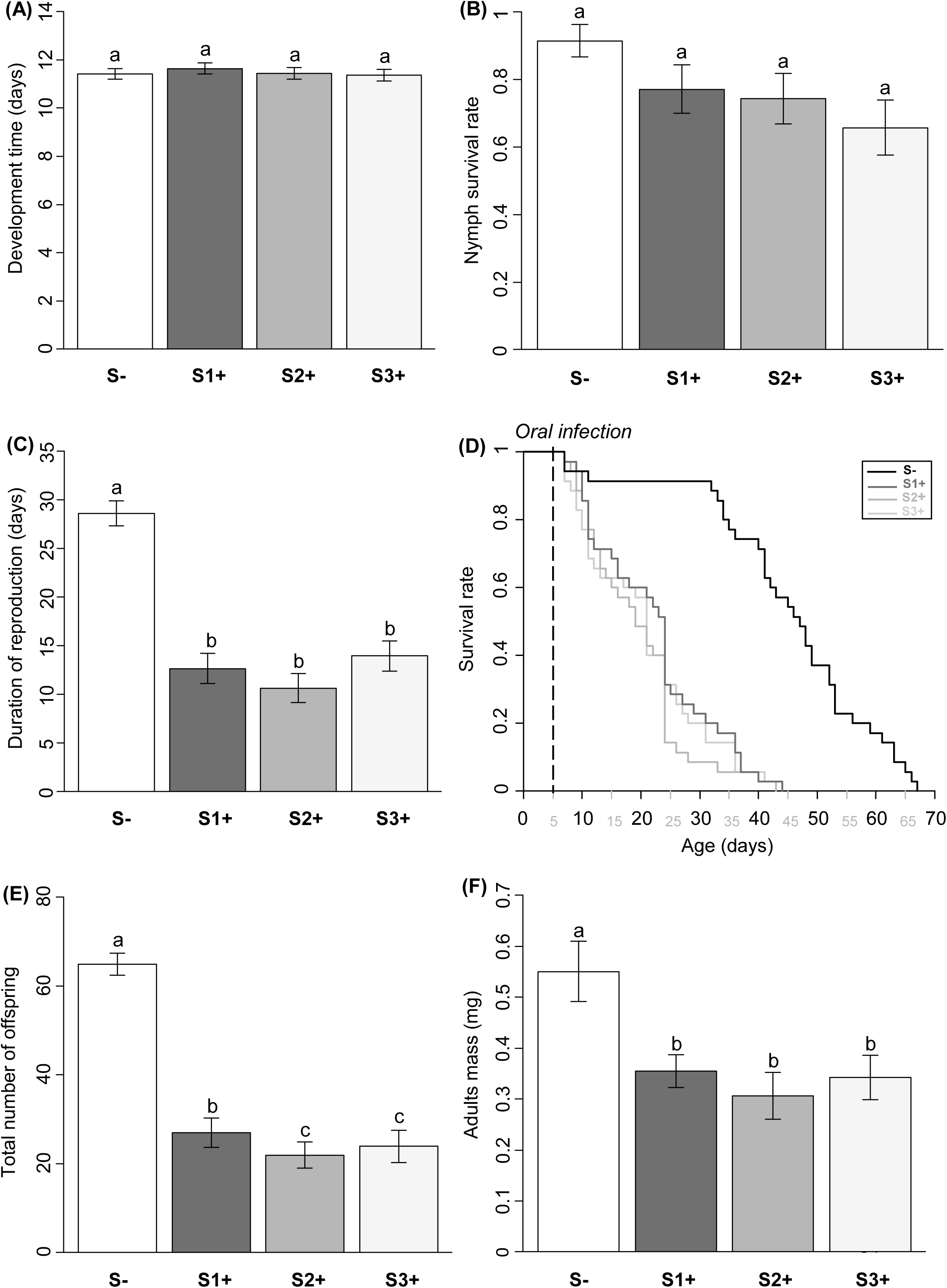
Effect of cultivable *S. symbiotica* on development time **(A)**, nymph survival rate **(B)**, duration of reproduction **(C)**, percentage of survivors **(D)**, total number of offspring **(E)** and Mass (mg) of adults **(F)** of the aphid *A. fabae.* Four host treatments were used: aphids uninfected with cultivable *S. symbiotica* (S-), aphids infected with cultivable *S. symbiotica* (S1 +: strain CWBI-2.3^T^), (S2 +: strain 24.1) and (S3 +: strain Apa8 A1). N=35. Error bars depict the standard error. Different letters show significant differences.

**TABLE 1.**
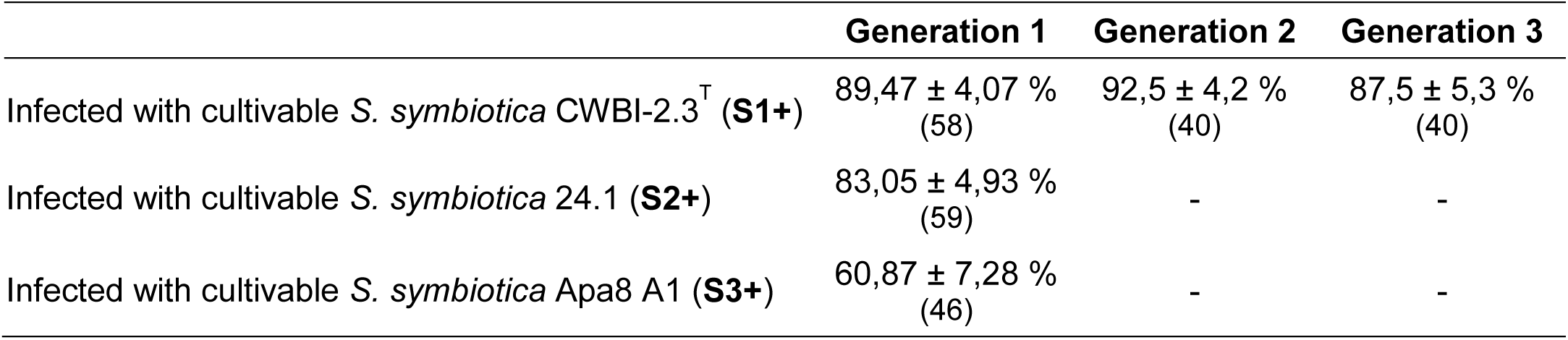
Effect of cultivable *S. symbiotica* on the survival rate of the aphid *A. fabae. **β**:* estimated regression coefficient of the Cox’s proportional hazard model, **exp (*β*)** hazard ratio, **SE (*β*)**: coefficient standard errors, ***z:*** z-test value and ***p***: significance of the coefficient.

### Protective phenotype associated with cultivable *S. symbiotica*

The cultivable *S. symbiotica* **S1+** strain had a significant negative effect on the parasitism rate (i.e. the number of parasitized aphids that become mummies) of *A. fabae* following *A. colemani* parasitoid attack (GLMM, χ^2^ = 28.48, df = 1, p < 0.001). The formation of mummies was lowered by about 40 % for infected aphids by cultivable *S. symbiotica* (29 ± 3 %) compared to uninfected aphids (71 ± 3 %) (Fig. 3A). The unmummified aphids were found either alive or “dead” (generally not found on the plant). The cultivable *S. symbiotica* had a significant effect on the death of attacked aphids (GLMM, χ^2^ = 21.21, df = 1, p< 0.001), with 47 ± 3 *%* of “dead” infected aphids contrary to 21 ± 3 % of uninfected aphids. Nevertheless, the rate of alive aphids was significantly more important to the infected aphids (24 ± 3 %) contrary to uninfected aphids (8 ± 2 %) (GLMM, χ^2^ = 8.07, df = 1, p = 0.0045). The bacterial strain also had a significant negative effect on the emergence rate (i.e. the number of adult parasitoids that emerged from mummies/total number of mummies) of parasitoids from *A. fabae* (GLMM, χ^2^ = 18.78, df = 1, p < 0.001), and on the weight of emerged parasitoids (LMM, χ^2^ = 5.76, df = 1, p = 0.016). A reduction of about 35% in emergence rate of parasitoids from infected aphids (58 ± 6 %) was observed compared to parasitoids from uninfected aphids (94 ± 2 %) (Fig. 3B). After dissection of the non-emerged mummies, the presence of a dead parasitoid larva was systematically observed. The weight of emerged parasitoids from infected aphids was lower compared to parasitoids from uninfected aphids (Fig. 3C).

**FIG 3.**
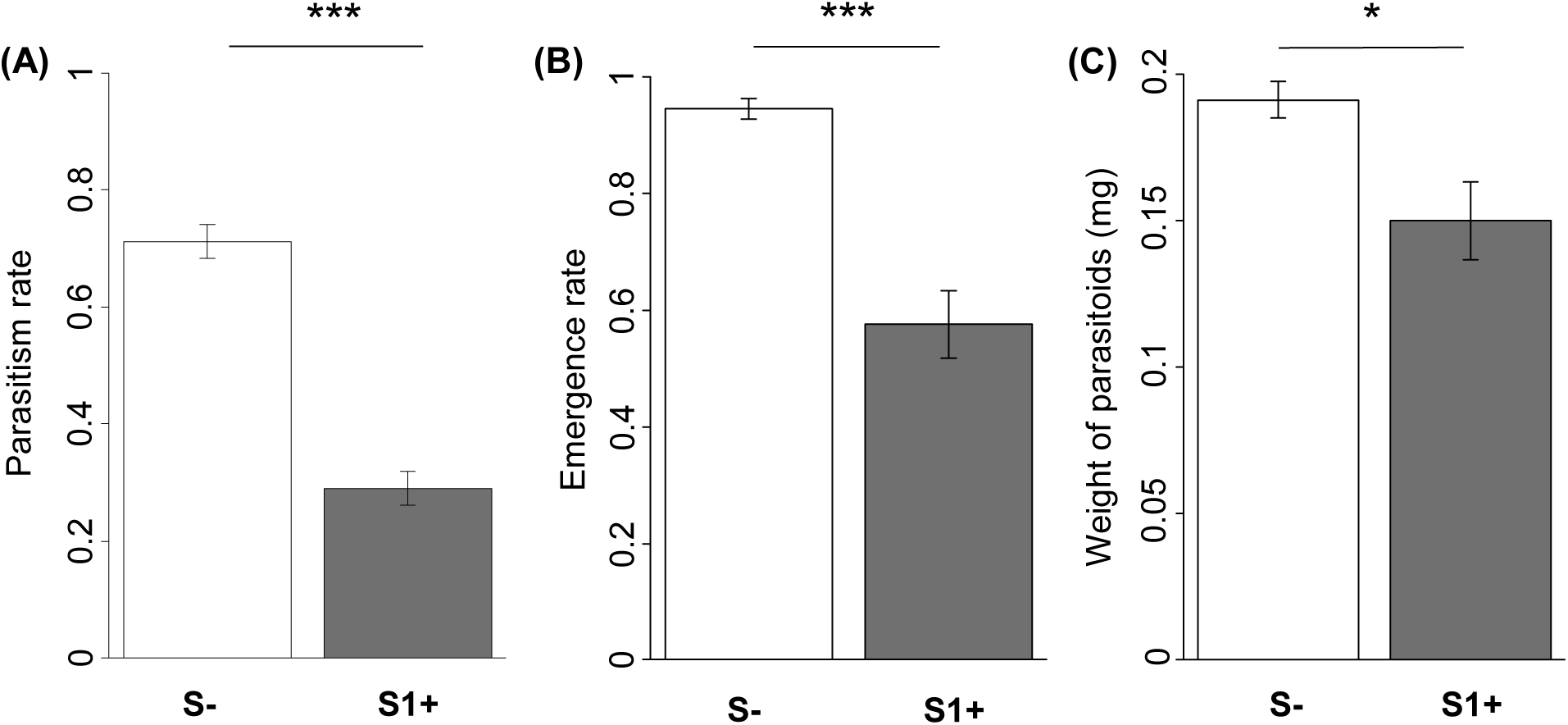
Effect of cultivable *S. symbiotica* on parasitism rate of *A. fabae* following *A. colemani* attack **(A)**, emergence rate of *A. colemani* from *A. fabae* **(B)** and weight (mg) of emerged parasitoids **(C)**. The parasitism rate is the number of parasitized aphids that become mummies and the emergence rate is the number of adult parasitoids that emerged from mummies. Two host treatments were used: aphids uninfected with cultivable *S. symbiotica* (S-) and aphids infected with cultivable *S. symbiotica* (S1 +: strain CWBI-2.3^T^). Error bars depict the standard error. *Stars* show significant differences (*: p<0.05 and ***: p<0.001).

### Cultivable *S. symbiotica* transmission mode

A high transmission rate of cultivable *S. symbiotica* was observed in the first aphid generation (Table 2), but significantly varied between strains (GLM, χ^2^ = 12.83, df = 2, p = 0.0016, Table 2). The transmission rate of **S1+** and **S2+** strains was higher compared to the **S3+** strain (Table 2). A high transmission rate of *S. symbiotica* **S1+** was also observed in the second and the third generation and the transmission over three generations was significantly similar (GLM, χ^2^ = 0.57, df = 2, p = 0.75, Table 2). For the other experiment, when aphids were taken off the plants immediately after birth (from adults infected by the **S1+** strain), none were infected with the bacteria, while seven out of ten offspring were infected with the bacteria when they remained on the plant with infected adults. The transmission was thus extracellular because required regular contacts between mother and offspring. We further detected the cultivable *S. symbiotica* **S1+** strain in honeydew samples collected from all thirteen aphids that were artificially infected in the digestive tract. These bacteria developed on the medium, indicating that living bacteria were transferred from mother to offspring.

**TABLE 2.**
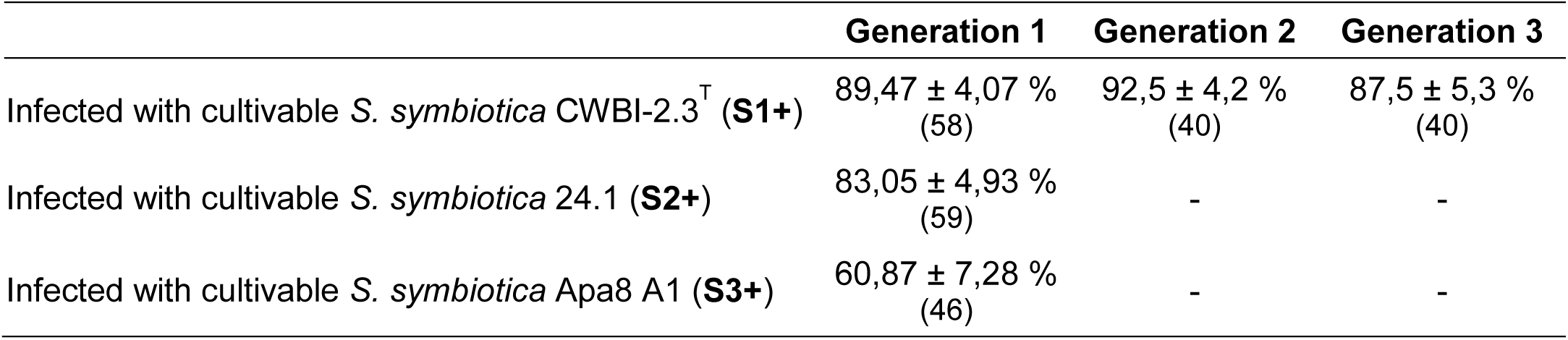
Percentage of transmission (± standard error) of cultivable *S. symbiotica* in first, second and third generation of *A. fabae.* Numbers in parentheses depict the number of individuals. Hyphens represent missing data.

### Cultivable *S. symbiotica* phylogeny

Phylogenetic analysis of concatenated sequences of *accD, gyrB, murE* and *recJ* genes revealed three distinct clades. Clade A was associated with the co-obligate endosymbiont *S. symbiotica* strains from *C. cedri* and *T. salignus* aphids (subfamily Lachninae, Fig. 5) which exhibit a long-term co-evolutionary history with their host. Clade B consisted of the co-obligate endosymbiont *S. symbiotica* strain from *C. tujafilina* and the facultative endosymbiont *S. symbiotica* strain from *A. pisum* (Fig. 5). Clade C included our three cultivable *S. symbiotica* strains (Fig. 5). Moreover, clades B and C were more closely related to each other than to clade A. We noticed that our cultivable strains are evolutionarily closer to their common ancestor than the co-obligate endosymbionts (clade A), but more distantly related than the *Serratia* species living independently of hosts (*S. marcescens* and *S. proteamaculans*). The cultivable strains are evolutionarily close to the clade of facultative endosymbionts.

**FIG 5.**
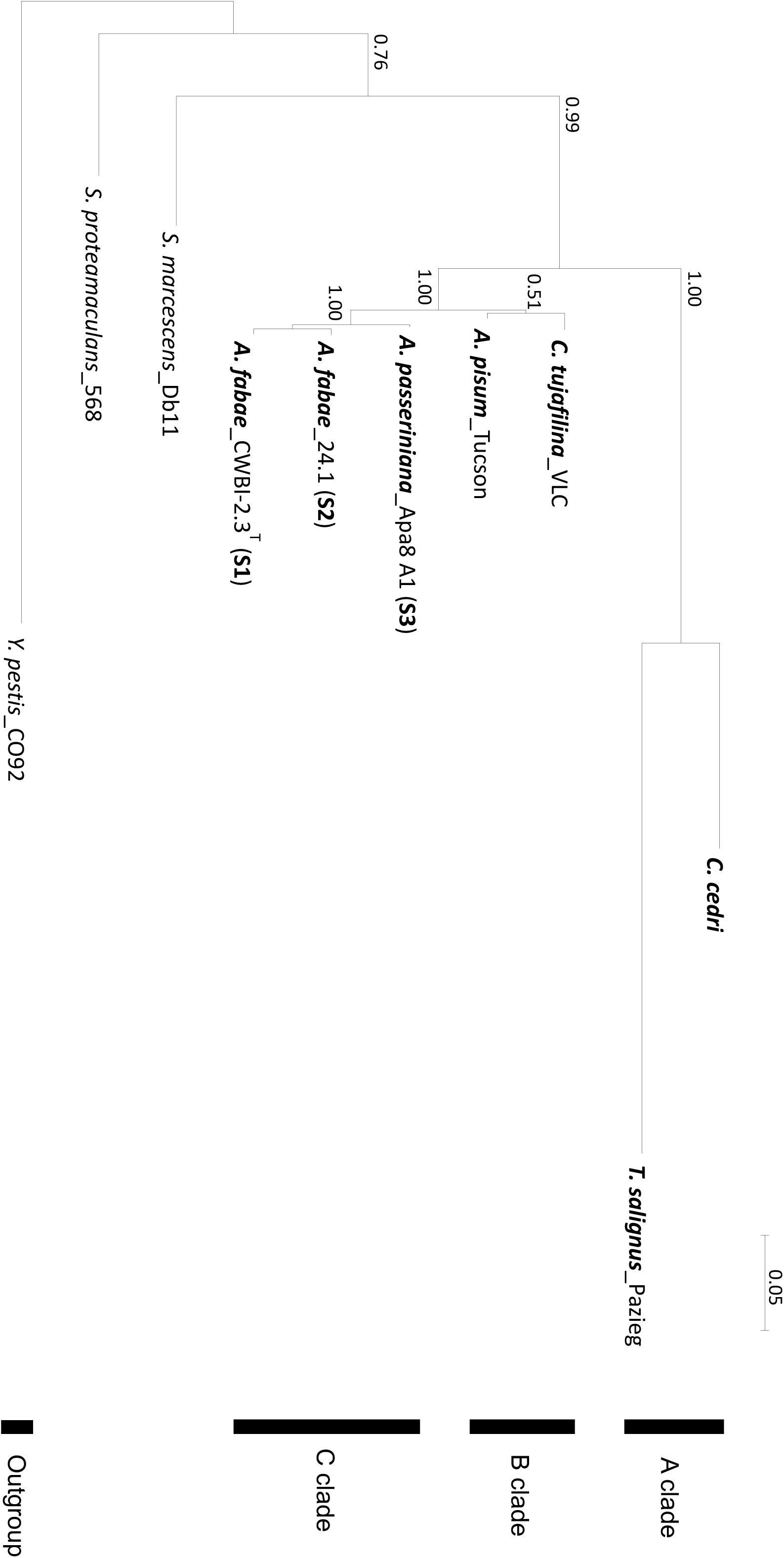
Phylogeny of *S. symbiotica* strains from different aphid species based on concatenated sequences of genes *accD*, *gyrB, murE* and *recJ* and constructed using Bayesian inferences. Name labels represent the aphid host of *S. symbiotica* followed by the *S. symbiotica* strains. *Serratia marcescens* and *Serratia proteamaculans* correspond to *Serratia* species living independently of hosts. *Yersinia pestis* was used as outgroup. Branch labels indicate marginal posterior probabilities.

## Discussion

The cultivable strains were isolated from *Aphis* species but were never clearly localized in their original hosts (31, 32). Nevertheless, a new field study revealed that these cultivable strains belong to the same clade that other *S. symbiotica* strains residing in the gut of field-collected aphids, mostly associated with *Aphis* genus (33). The bacteria *S. symbiotica* have typically been referred to as strict endosymbiont but the discovery of cultivable strains that are potentially capable of growing outside aphid hosts shows a different aspect of this symbiont species and raises the questions of the biological effects of such strains in aphid hosts, as well as the nature and the durability of these associations.

We found that the oral infection was reliable and aphid gut was suitable for harboring cultivable *S. symbiotica* bacteria. This bacterium multiplied and spread throughout the whole digestive tract, with bacterial densities increasing exponentially with aphid age (similar of aphid endosymbionts) (30, 38, 39). This infection seems similar to infections found in field-collected aphids (33). The bacteria do not seem to be found in specialized structures such as with the *Burkholderia* gut symbiont of the bean bug *Riptortus pedestris* (40) and *Sodalis glossinidius* of *Glossina morsitans* (41). Indeed, due to the plant-sap diet, aphids have very distinct guts that are unlikely to have crypts (42). To evolve an association with a host, bacteria must be able to pass through the host’s defense mechanisms and have a battery of molecular tools to facilitate the infection (13). Our results suggest that the cultivable bacterium exhibits suitable mechanisms to invade the aphids’ gut, which is consistent with a study that provide evidence that the bacterium is well-armed to develop in the aphids’ gut (36). Another study showed that the cultivable *S. symbiotica* is not perceived by the host's immune system as an antagonist contrary to the pathogen *Serratia marcescens* (43), allowing the bacterium to multiply and persist in the new host.

Once infection has occurred, the aphid host must not succumb to the bacteria to allow the initiation of a long standing interaction. Our results demonstrated that cultivable *S. symbiotica* gut infections have a negative effect on some *A. fabae* host life history traits in the absence of environmental stressors. These are similar with previous studies that underlined the induced cost of facultative endosymbionts for aphids (23, 25, 44, 45). There may be several potential causes of virulence of the cultivable bacteria. The important proliferation of the bacteria may explain the fitness cost observed. Indeed, a correlation between virulence and bacterial density has been documented for *Wolbachia* infecting *Drosophila melanogaster*, but showing that the density and fitness costs associated with infection rapidly decline over generations (46, 47). Indeed, it has been shown that the origin of mutualistic symbiosis in nature generally occurs by acquiring free-living bacteria initially pathogens, which have progressively lost their antagonistic character during the processes of evolutionary degeneration (12, 17). Here, cultivable strains have conserved an array of molecular tools facilitating the invasion in new hosts such as virulence factors (34, 36). The bacteria therefore have the potential to be harmful, but the cost associated with its establishment can ameliorate in subsequent generation. In addition, the magnitude of costs can depend on the combination of aphid genotype and symbiont strain (45), thus harboring a non-adapted bacteria can also result in negative fitness consequences for the host (48). These findings demonstrate that our cultivable strains have a potential to be virulent in the first step of infection. However, if the hosts survive and the bacteria confer beneficial effects to their hosts, the symbiosis could stabilize and persist.

We showed that the aphids infected by cultivable *S. symbiotica* in their gut exhibited a great resistance to parasitism by *A. colemani* compared to uninfected aphids in term of mummy formation, emergence rate and weight of adult parasitoids. Previous studies already showed the protective effect of *S. symbiotica* endosymbionts against parasitoids (29, 49), but not so important. The mechanism by which *S. symbiontica* protect their aphid hosts is still not well understood (4), but studies showed that the protection associated with another aphid symbionts, *Hamiltonella defensa* is due to its interaction with APSE phages (30, 50). In our case, the genome of the cultivable strain (CWBI-2.3^T^) does not contain APSE phages (Renoz *et al.*, in prep). The mechanisms of protection are likely to be different. The endosymbionts occur in the aphid hemolymph and the parasitoid egg is directly exposed to them (4) while the cultivable bacteria are dwelling in the gut of aphids. One hypothesis could be that the protection may be due to the effects of the bacteria on aphids causing a less favorable environment for the development of parasitoid larvae. This opens the question of the protective effect observed in other aphid endosymbionts that is not really explained. The fitness cost of symbionts on aphid host could partly explain the mechanism of protection against parasitoids. Other putative beneficial effects associated with cultivable *S. symbiotica* will have to be studied to clarify their implication in the ecology and evolution of aphids. They could protect their hosts against other stresses (51, 52), but they also could be a contributor to the nutrition of their hosts because of their conservation of a large repertoire of genes related to metabolism (34). Thereby, our results highlight that the bacteria can be both parasites and mutualists depending on the context. They can confer immediate benefits to their new hosts, which may offset costs of infection in this context facilitating the establishment of a stable mutualistic interaction and explaining the presence of these strains in the nature.

Vertical symbiont transmission from mother to offspring is one of the most important processes for the establishment and maintenance of intimate host-symbiont associations (53), although stable symbiosis without vertical transfer are also found in nature (40, 54). Here, the occurrence of transmission may be questioned, because gut bacteria generally lack a reliable transmission pathway (55). The transmission rate of cultivable *S. symbiotica* strains over three generations of aphid remained high, although it was lower than the rates of vertical transmission observed for facultative endosymbionts of aphids (4). We showed that transmission of the cultivable strains requires contact with the infected mothers otherwise offspring are not infected, implying that nymphs acquire bacteria after birth (56) and that the bacteria are not transmitted through the same mechanism used by maternally-transmitted symbionts that live within the body cavity. A study showed that honeydew can mediate transfers of symbionts in aphids (57), where bacteria on plant surfaces are picked up by offspring. This could be the case for the cultivable *S. symbiotica* because bacteria were detected in honeydew samples of aphids artificially infected in the gut. However, it is also possible that aphids pick them up through circulation of the bacteria in the host plant. This transmission mechanism through the phloem of the plant has already been observed in some insect systems, such as *Rickettsia* with *Bemisia tabaci* (16) and *Cardinium* with leafhoppers (58). Further studies are thus needed to analyze this tripartite interaction to know if the plant could be a way of cultivable *S. symbiotica* transfer. The possible transmission of the cultivable bacteria suggests a durability of the novel association but further tests will have to be realized to complete our knowledges.

The wide diversity of *S. symbiotica* strains varying in their degree of reliance on hosts suggests that this symbiont species is embedded in diverse evolutionary paths (36). Our phylogenetic investigation supports this suggestion. Among this gradient, the cultivable *S. symbiotica* strains have intermediate genome characteristics between a free-living bacterium and a facultative endosymbiont (35), suggesting that the cultivable strains are involved in a nascent stage of symbiosis with aphids and would have diverged little from their free-living counterparts. Their potential localization and independence with respect to their hosts propose that a direct acquisition by the environment is not excluded. Through phylogenetic and evolutionary analyses, it is well established that endosymbionts have evolved from free-living ancestors residing in the environment but which have gradually domesticated by their hosts (12, 59). According to personal observations, the *S. symbiotica* could be found thriving in the aphid's natural environment. We thus hypothesized that the endosymbiont *S. symbiotica* strains evolved from free-living precursors via the oral entry route and that environmental generated infections could represent a reservoir of new beneficial traits for hosts creating a pathway towards new symbiotic associations. Gut associated cultivable strains could then cross the epithelial barrier of the midgut to infect the host’s hemolymph via transcytosis mechanism (60). However, we did not observed any red fluorescence in the hemolymph of aphids suggesting that cultivable bacteria require more time or were not capable of crossing the aphid gut epithelium. Moreover, artificial infection in the aphid body cavity via microinjection are also perform to determine whether the strains are able to live in the hemolymph and no infection was maintained over time. However, the establishment of aphid symbionts can be dependent on host genotype (61), thus the maintenance could be different for other associations. Alternatively, cultivable *S. symbiotica* could be strictly adapted to the gut of aphids and represent a new kind of symbionts. The bacteria could be pre-adapted to develop in aphid gut and form a stable symbiotic association or they may not be on evolutionary trajectory towards greater intimacy.

In conclusion, the identification of strains with free-living capability is very interesting because it allows the experimental traceability of the association and may contribute to our understanding of the mechanisms that shape symbiosis in insects. Our study help to clarify the nature of symbiosis in aphids, revealing that cultivable *S. symbiotica* strains have the ability to rapidly infect the aphid gut and seem to be transmitted over generations through an external transmission mechanism. The bacteria also induced a fitness costs on their new hosts when colonizing their entire digestive tract. However, they offered immediate fitness benefits under environmental stress. More studies are needed to support our findings and answer questions: What is the origin of these strains? What is their prevalence in natural aphid populations? Can they have a free-living capacity? Or are they strictly associated with the aphid gut? Are they originally present in the environment and capable of transiting through plants? How are they perceived by the immune system of their new host? Are they capable of transfer across the aphid gut wall? Do they form a stable association with aphids? Are these strains a first step to become a vertically transmitted symbiont or are they an alternative way of being an aphid symbiont? Other studies in other insect-symbiont systems assessing the diversity of symbiotic associations are also needed, to help us enhance our perception of the evolution of symbiosis.

## Materials And Methods

### Insects and Bacterial strains

A single clone, A06–407, of *Aphis fabae* was originally collected from *Chenopodium album* in St. Margrethen (Switzerland) and provided by Christoph Vorburger (Eawag, Switzerland). This clone was found to be uninfected with any known facultative symbionts of aphids (62). Aphids were maintained through parthenogenetic reproduction on *Vicia faba* at 18°C under a long-day photoperiod (16L: 8D) and 65 ± 3 % of humidity. The aphid parasitoid, *Aphidius colemani* (Hymenoptera: Aphidiinae), used for the experiment was provided by Viridaxis SA, Belgium in April 2016. It is a generalist parasitoid which naturally attack *A. fabae* host (63). The *S. symbiotica* strain CWBI-2.3^T^ (**S1+**), isolated from a field-collected *A. fabae* (31, 34), was used in this study. In addition, we isolated two novel strains of *S. symbiotica* as described in (31, 32): the strain 24.1 (**S2+**) from a field-collected *A. fabae* and the strain Apa8 A1 (**S3+**) from a *Aphis passeriniana* collected in Tunisia. These strains, having a free-living capacity, were preserved in frozen stocks at -80°C and cultured at 20°C in 863 medium (1% yeast extract, 1% casein peptone, 1% glucose) as described in (31), not containing insect cell lines and FBS (fetal bovine serum) contrary to some aphid symbionts already cultured outside of aphids but with enriched medium that are designed for insect cell culture with insect cell lines and/or FBS (50, 64).

### Diagnostic PCR

To verify the integrity of the *A. fabae* clone before infection with bacterial strains, as well as the presence of *S. symbiotica* in aphids after the inoculation procedure, DNA from individual aphids was extracted by using the QIAamp tissue kit (Qiagen). PCR primers used for *S. symbiotica* detection (the three strains) were 16SA1 (AGAGTTTGATCMTGGCTCAG) and PASScmp (GCAATGTCTTATTAACACAT) (65). PCR reactions were performed in a final volume of 15 μl containing 1 μl of the template DNA lysate, 0.5 μM of each primer, 200-μM dNTP’s, 1X buffer and 0.625 unit of Taq DNA polymerase (Roche). PCR reaction conditions consisted of 35 cycles at 95 °C for 30 sec, 55 °C for 1.5 min and 72 °C for 1.5 min. Moreover, before the start of the experiments, to ensure clonal integrity, the aphid clone was diagnosed as uninfected with known facultative endosymbionts of aphids by diagnostic PCR (62).

### Oral infection

Oral infection of cultivable strains of *S. symbiotica* was performed by feeding aphid hosts on an artificial medium containing the desired strain to ensure the presence of the bacteria in the digestive tract (66), to mimic what is found in nature (33). Bacterial strains were first grown to an early log phase in 863 medium (31) on a gyratory shaker (160 rpm) at 20°C. When an optical density (OD) between 0.5 and 0.7 at 600 nm was reached during the growth phase, bacteria were centrifuged. Symbiont cells were then washed with sterile PBS (Sigma) and suspended in PBS to obtain an OD of 1 at 600 nm. To standardize-aphid individuals, adult females of *A. fabae* were left on young *Vicia faba* plants for 24 h to produce nymphs. After removal of the adult insects, the newborn nymphs were kept on the same plants for four days prior to infection experiments. Third-instar aphid nymphs were then fed on an artificial diet (67) for 24 h. One hundred μl of bacterial solution (sterile PBS for the control) was mixed with 20 ml aphid diet (approximately 10^6^ CFU/ml of diet for each strain; as found in (43, 66)).

### Histological observations

To visualize the infection process of cultivable *S. symbiotica* **S1+** strain, whole-mount fluorescence *in situ* hybridization (FISH) was performed, as previously described in (43, 68). At different times after the infection procedure (0, 2, 5, 10, 15, 20 and 30 days post-ingestion), sampled aphids were placed in acetone for preservation. The following oligonucleotide probes were used: Cy5-ApisP2a (CCTCTTTTGGGTAGATCC) targeting 16S rRNA of *B. aphidicola* and Cy3-PASSisR (CCCGACTTTATCGCTGGC) targeting 16S rRNA of *S. symbiotica.* Samples were observed under a Zeiss LSM 710 confocal microscope. Negative controls consisted of aphids not infected with *S. symbiotica*, with the two probes (Fig S1, Supporting information) and infected aphids with no-probe staining.

### Cultivable *S. symbiotica* density

The infection dynamic of a cultivable *S. symbiotica* **S1+** strain was measured by TaqMan real-time quantitative PCR on an Applied Biosystems Step One Plus machine (Applied Biosystems), as previously described in (38). The development of *S. symbiotica* density relative to aphid growth was quantified in six replicates collected at different time points after bacterial administration: 0, 2, 5, 10, 15, 20 and 25 days after ingestion. To estimate *S. symbiotica* density, the copy number of *S. symbiotica’*s *dnaK* gene was quantified with the following primers and probe (30): forward TGGCGGGTGATGTGAAG, reverse CGGGATAGTGGTGTTTTTGG and probe ATTGAAACTATGGGCAGCGTGAT. As an index of aphid cell number, the copy number of *A.fabae’s EFla* gene was quantified using the following primers and probe (38): forward CAGCAGTTACATCAAGAAGATTGG, reverse CATGTTGTCTCCATTCCATCCAG and probe CCCAGCCGCTGTTGCTTTCGTTCC. Probes were modified with FAM as the 5′-terminal reporter dye and BHQ-1 as the 3′ terminal quencher dye. DNA extraction from whole aphid was conducted using a Qiagen DNeasy kit. QPCRs were performed in duplicate using a total volume of 25 μL containing 5 μL of template DNA. The PCR reaction mixture included 12.5-μl Maxima Probe/ROX qPCR Master Mix, 6.7-μl ddH_2_O, 0.3 μl of each primer (10 nM), and 0.2 μl of Probe. The cycling conditions were: 10 min activation at 95 °C followed by 40 cycles at 95 °C for 15 s, at 60 °C for 30 s, and 72 °C for 30 s. Gene copy number was estimated from the standard curve generated by a 10-fold series of dilutions of genomic DNA at concentrations of 10^2^, 10^3^, 10^4^, 10^5^, 10^6^, 10^7^ and 10^8^ copies/μl.

### Life-history traits measurement

To measure potential fitness costs of the infection with cultivable *S. symbiotica* strains, third-instar aphid nymphs were standardized and fed on an artificial diet (67) with bacterial solution (or sterile PBS solution for the control) for 24h. Four different host conditions were tested: uninfected aphids (control), aphids infected with **S1+** strain, infected with **S2+** strain and infected with **S3+** strain. After oral ingestion, the aphids (35 wingless third-instar nymphs per conditions) were individually placed on young *V. faba* plants and six life history traits were analyzed. Each third-instar aphid nymph was observed daily until death to measure nymph survival rate, development time (i.e. time from birth to the onset of reproduction), total fecundity (i.e. total number of offspring produced by each aphid until death), duration of reproduction (i.e. time from the first to last offspring production), and lifespan. To estimate the body size, the fresh mass of a set of different adult individuals (between 9 and 18) was measured at 10 days of age using a Mettler MT5 microbalance (Mettler Toledo, Switzerland).

### Parasitism assays

To measure potential fitness benefits of the infection with cultivable *S. symbiotica* strain, eighteen third-instar aphids were introduced in a 4-cm diameter glass Petri dish containing a *V. faba* leaf, after oral ingestion by the aphids. Two different aphid treatments were tested: uninfected aphids (control) and aphids infected with **S1+** strain. A single parasitoid female was introduced into the Petri dish, with 2 or 3 days-old, mated and fed. Before the experiment, each parasitoid was exposed to one third-instar aphid for oviposition experience. The aphids were removed from the Petri dish once an ovipositor insertion was observed (70). After 30 minutes, attacked aphids were transferred onto a *V. faba* plant and twelve days later, they were inspected to measure the parasitism rate, estimated by dividing the total number of aphid mummied (i.e. dead aphids containing a developing parasitoid) by the total number of attacked aphids. The rates of live and dead aphids were also measured. Seven days later, the mummies were inspected to measure the emergence rate of parasitoids, estimated by dividing the number of adults emerging from mummies among the total number of mummies. Each emerged parasitoid was weighed using a Mettler MT5 microbalance (Mettler Toledo, Switzerland) and the non-emerged mummies were dissected. We performed sixteen experimental replicates for aphids infected and fifteen for uninfected aphids.

#### Symbiont transmission mode

##### (i) Transfer across host generations

Offspring (10 days-old) from the first generation of adult aphids infected artificially at the gut level by cultivable *S. symbiotica*, were collected and subjected to diagnostic PCR for all strains (**S1+** (N=58); **S2+** (N=59) and **S3+** (N=46)). A single diagnostic PCR was used for all three strains. Offspring from the second (N=40) and third (N=40) generations were also collected for only **S1+** strain and subjected to diagnostic PCR.

To investigate how the transmission of cultivable *S. symbiotica* (**S1+**) was carried out, another experiment was performed. Five infected aphids (by **S1+** strain) were placed on a *V. faba* plant. Ten offspring were immediately taken off the plant when born and placed on a new plant to verify if the transmission of the bacteria was carried out after birth or required regular contacts with infected adults. Ten other offspring were left on the plant with infected adults. The offspring (10 days-old) were then collected and subjected to diagnostic PCR. Before each DNA extraction, aphids were rinsed with ethanol 99%, bleach then pure water.

##### (ii) Honeydew inspection

Honeydew from thirteen adult aphids infected artificially at the gut level by a cultivable *S. symbiotica* (strain **S1+**, 9 days post-infection) was collected and subjected to diagnostic PCR. Aphids were placed on an artificial diet for one day to harvest the honeydew. Separate honeydew samples were then placed on 863 medium at 20°C to observe whether the *S. symbiotica* were alive in the honeydew of the aphids. Sequencing was carried out to verify the identity of recovered *S. symbiotica* strain.

### Phylogenetic analyses

A multi-locus sequence typing (MLST) was performed for *S. symbiotica* using the partial sequence of four housekeeping genes *accD, gyrB, murE* and *recJ* (Table S1, Supporting information) (71). PCR amplification was realized on DNA samples of all three cultivable *S. symbiotica* strains under the following conditions: initial denaturation at 94°C for 5 min; 35 cycles of denaturation at 94°C for 30 s, annealing at 60-64°C depending on primers for 30s, extension at 72°C for 1 min; a final extension at 72°C for 5 min (33). Amplicons were purified and sequenced in both directions (Macrogen Inc., Amsterdam). Sequences obtained were cleaned and aligned using Geneious^®^ v9.1.5 (72). Phylogenetic associations were analyzed for sequences from the three cultivable *S. symbiotica* strains, sequences from *S. symbiotica* whose genomes were already known (28, 73), as well as sequences from *Serratia marcescens* Db11 and *Serratia proteamaculans* 568 which correspond to *Serratia* species living independently of hosts (74, 75) and a sequence from *Yersinia pestis* CO92 (outgroup). All sequences are available in GenBank (Table S2-S4, Supporting information). SeaView v4.6.1 was used to align sequences and GBlocks v0.91b (76) to remove poorly aligned positions and divergent regions of DNA alignments. PartitionFinder v1.1.0 (77) was used to determine the evolutionary model of sequences for each gene: K80 model for *accD* gene, HKY model (1,2) and K80 model (3) for *gyrB* gene, K80+G model (1,2) and K80 model (3) for *murE* gene and K80 model (1,2) and HKY+G model (3) for *recJ* model. Phylogenetic analysis was performed by Bayesian inference methods with MrBayes v3.2.6 (10^6^ generations) (78).

### Statistical analyses

The effect of cultivable *S. symbiotica* strains were tested on the following dependent variables: nymph survival rate, development time, adult fresh mass, survival rate, total number of offspring and duration of reproduction of aphids. Survival data were analyzed with a proportional hazards regression (79) and visualized by computing the Kaplan-Meier survival functions for the different infections (80). Mass of aphids was compared with general linear model, after verification of normality of the data. All other variables were analyzed using generalized linear models with a Binomial error structure and a logit-link function for nymph survival rate, a Gamma error structure and an inverse-link function for development time and duration of reproduction, and a Poisson error structure and log-link function for a total number of offspring. Analyses of parasitism and emergence rates, as well as, the rates of live and dead attacked aphids were performed by fitting generalized linear mixed models by assuming a Binomial error structure and a logit-link function, while the weight of parasitoids was analyzed using general linear mixed model, after validation of normality of the data, testing for the effect of cultivable *S. symbiotica.* The parasitoid female was considered as a random factor in these statistical modeling because one parasitoid female attacked several aphids. We further analyzed the vertical transmission rate using generalized linear model with a Binomial error structure and a logit-link function, testing for the effect of different cultivable *S. symbiotica* strains and generations. Statistical analyses were performed using the software R version 3.0.1 (R Development Core team, 2014), using the *survival* package for survival analyses (81), the *lme4* package for linear mixed models and the *GrapheR* package for graphics.

## Data availability

DNA sequences generated during this study have been submitted to GenBank and accession numbers are available in supporting information (S2, S3 & S4 Tables).

## Acknowledgments

We thank Christoph Vorburger who supplied the A06–407 clone *Aphis fabae* used in our experiments. We thank Charles Hachez for FISH assistance, Gwennaël Bataille for his help on the phylogenetic analysis, as well as Huma Khalil for her help on the parasitism experiment, Sarah Becker for qPCR assistance and the team of Nicolas Schtickzelle for technical facilities. We are very grateful to Bertanne Visser, Florence Hecq and Guillaume Le Goff for their helpful comments and corrections on the manuscript.

This work was supported by the Fonds de la Recherche Scientifique (FNRS) through a Fonds pour la Formation à la Recherche dans l’Industrie et dans l’Agriculture (FRIA). This paper is publication BRC 395 of the Biodiversity Research Center (Universtité catholique de Louvain).

